# The radial expression of dorsal-ventral patterning genes in placozoans, *Trichoplax adhaerens,* argues for an oral-aboral axis

**DOI:** 10.1101/345777

**Authors:** Timothy Q. DuBuc, Yuriy Bobkov, Joseph Ryan, Mark Q. Martindale

## Abstract

The placozoans are a morphologically simplistic group of marine animals found globally in tropical and subtropical environments. They consist of a single named species, *Trichoplax adhaerens* and have roughly six morphologically distinct cell types. With a sequenced genome, a limited number of cell-types and a simple flattened morphology, *Trichoplax* is an ideal model organism to understand cellular dynamics and tissue patterning in the first animals. Using new approaches for identification of gene expression patterns this research looks at the relationship of Chordin/TgfB signaling and the axial patterning system of Placozoa. Our results suggest that placozoans have an oral-aboral axis similar to cnidarians and that the parahoxozoan ancestor (common ancestor of Placozoa and Cnidaria) was likely radially symmetric.

## Introduction

The phylum Placozoa is an unusual group of animals currently represented by a single described species *Trichoplax adhaerens.* Placozoans are marine animals found on biofilm surfaces around tropical and subtropical environments [1, 2]. They are one of four lineages that separated from the main animal trunk prior to the last common bilaterian ancestor, along with Cnidaria (e.g. corals, sea anenomes, jellyfish and *Hydra*), Porifera (sponges) and Ctenophora (comb jellies). Most phylogenetic analyses places them as the closest outgroup to the cnidarian-bilaterian ancestor[3–7] (Figure 1a), yet their simple morphology have led some to speculate that the modern day placozoan “body plan” is similar to, and descended directly from, that of the last common ancestor of all animals[8, 9]. They consist of a flagelated upper and lower epithelial layer, thought by some to correspond to the dorsal and ventral axis of Bilateria [10–12] (Figure 1b-c). Total genome sequencing studies have shown that placozoans have a rather complete developmental toolkit for body patterning relative to bilaterians [3, 13, 14], therefore, it is confounding why these mysterious creatures exibit such a small and seemingly “simple” body plan.

**Figure 1.**
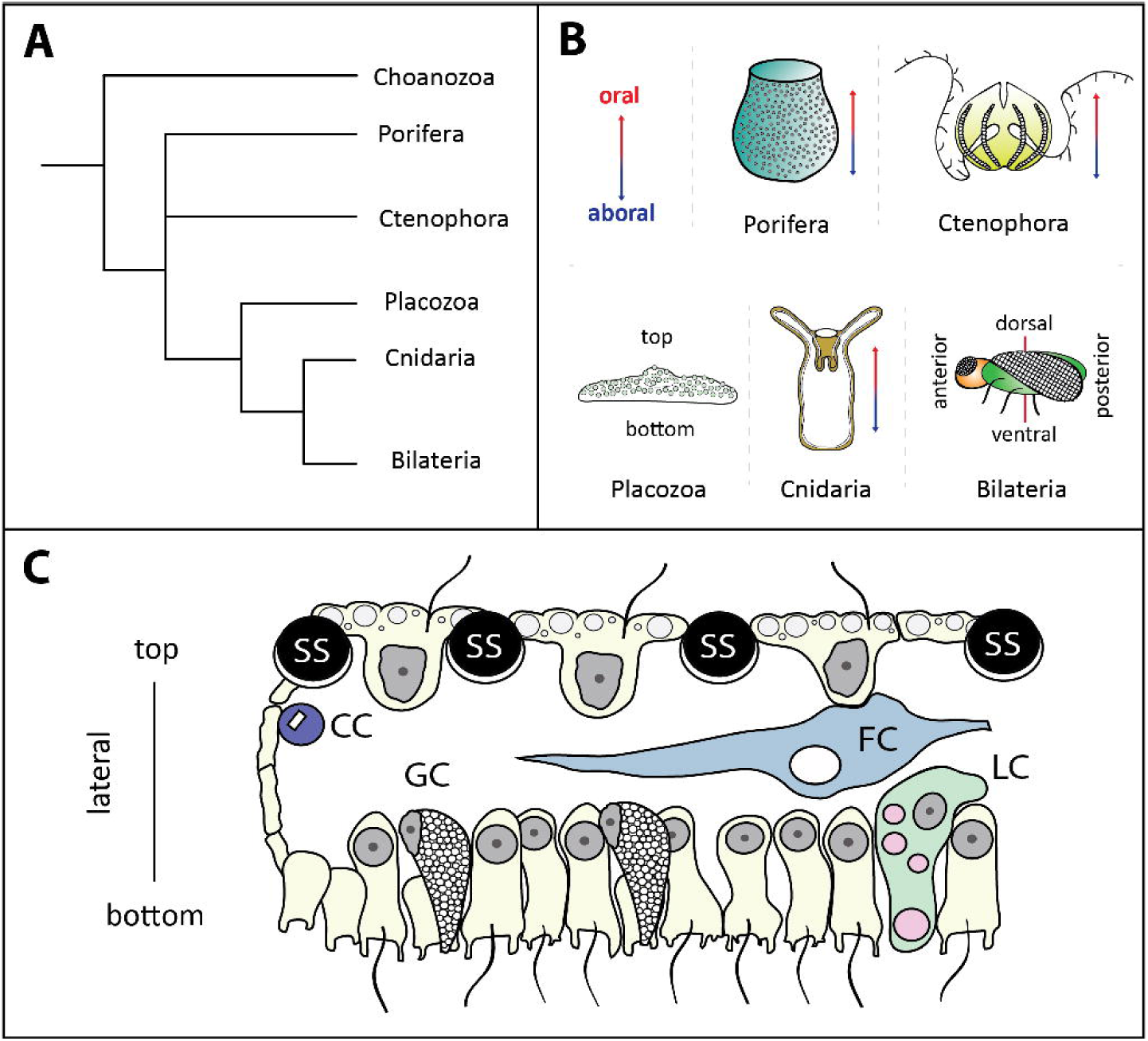
Phylogenetic relationships and body axes of the five major animal lineages. A) Phylogenetic relationship of animals consistent with the vast majority of animal phylogenomic studies (including [3–7]). B) Body axes of metazoan lineages. It has been speculated that the top-bottom axis of Placozoa is homologous to the dorsal-ventral axis of Bilateria. C) Schematic diagram of the top, middle and bottom tissue layers of placozoans[12]. The upper layer consists of a thin ciliated epithelial layer, with interspersed shiny spherical cells (SS) thought to be specific to Placozoa. The middle layer has fiber cells (FC) and crystal cells (CC) of unknown functionality. The bottom layer is a thick ciliated epithelial layer interspersed with gland cells (GC) and lipophil cells (LC) used for digestive function.

Karl Grell first described the lower epithelial layer composed of flagellated, cylindrical cells and scattered aflagellated gland cells [12]. The upper layer consists of ciliated epithelial cells interspersed with autofluorescent shiny sphericals thought to be used in predator defense (Figure 1c) [15]. Between the upper and lower layers, numerous small, ovoid cells (theorised to be stem cells [16]) are positioned at the margin along the lateral edge (Figure 1c); these have been described in detail by ultrastructural analyses [17]. Also in this internal layer, there are branching fiber cells, the cell bodies of secreting lipophil cells and crystal cells, which each contain a birefringent crystal [18, 19]. The lateral edge of the adult animal is thought to have neuro-sensory functionality based on localization of the neuropeptide RFamide protein in dispersed cells along the edge [20].

Interestingly, unlike in most other model systems, gene expression analyses have raised more questions than they have supplied answers towards understanding the morphology of *Trichoplax.* Only a handful of genes have been described by *in situ* hybridization in *Trichoplax.* A number of these genes (*Brachyury, PaxB, Secp1, Trox-2*) were shown as being expressed along a ring around the perimeter of the animal, within cells of middle tissue layer [16, 21–23]. The homeobox gene *Tbx2/3* was shown to be expressed in both upper and lower tissue layers [21], and expression of another homeobox gene, *Not,* was reported in the bottom layer in folded tissue along the lateral edge[22]. Actin expression has been reported both throughout[16] and within a patch in the middle *Trichoplax* [23]. Due to the large reoccurrance of a single spatial gene expression pattern, a ring near the lateral edge of the animal, it has been difficult to use gene expression to compare cell types and groups of cells between *Trichoplax* and other animals. This study was partially motivated by this constraint caused by this odd set of spatial coincidences, especially in the context of genes known to pattern the secondary directive axis (dorsal/ventral) axis in most other animals.

Herein, we have developed and applied an improved RNA *in situ* hybridization technique for *Trichoplax.* With this protocol in hand, we have determined the spatial gene expression patterns for key genes involved in dorsal/ventral patterning in bilaterians. These patterns reveal that *Trichoplax* exhibit dynamic spatial gene expression patterns, and provide new evidence towards understanding the relationship between the body axes of Placozoa and other animals. We anticipate that our new protocol will empower a more complete genetic understanding underpinning the biology of this fascinating animal.

## Results

### Living on a biofilm

Placozoans cohabit hostile marine environments with diverse unicellular organisms, inverbrate predators and high-energy tides and waves. In cultures, they are often sustained on a single species of algae, supplemented with algal growth media and live in a motionless environment [24–26]. Individual animals can be removed from the biofilm surface through gentle pipette propulsion and can be transfered to new dishes (Figure 2a). When collecting individuals, one side of the animal often appears to adhere to the surface with greater adhesion, regardless of the position of the pipette (Figure 2b). The morphological evidence does not suggest any assymetries, but there may exist a zone of attachment along the lateral edge of the animal, although futher study is warrented.

**Figure 2.**
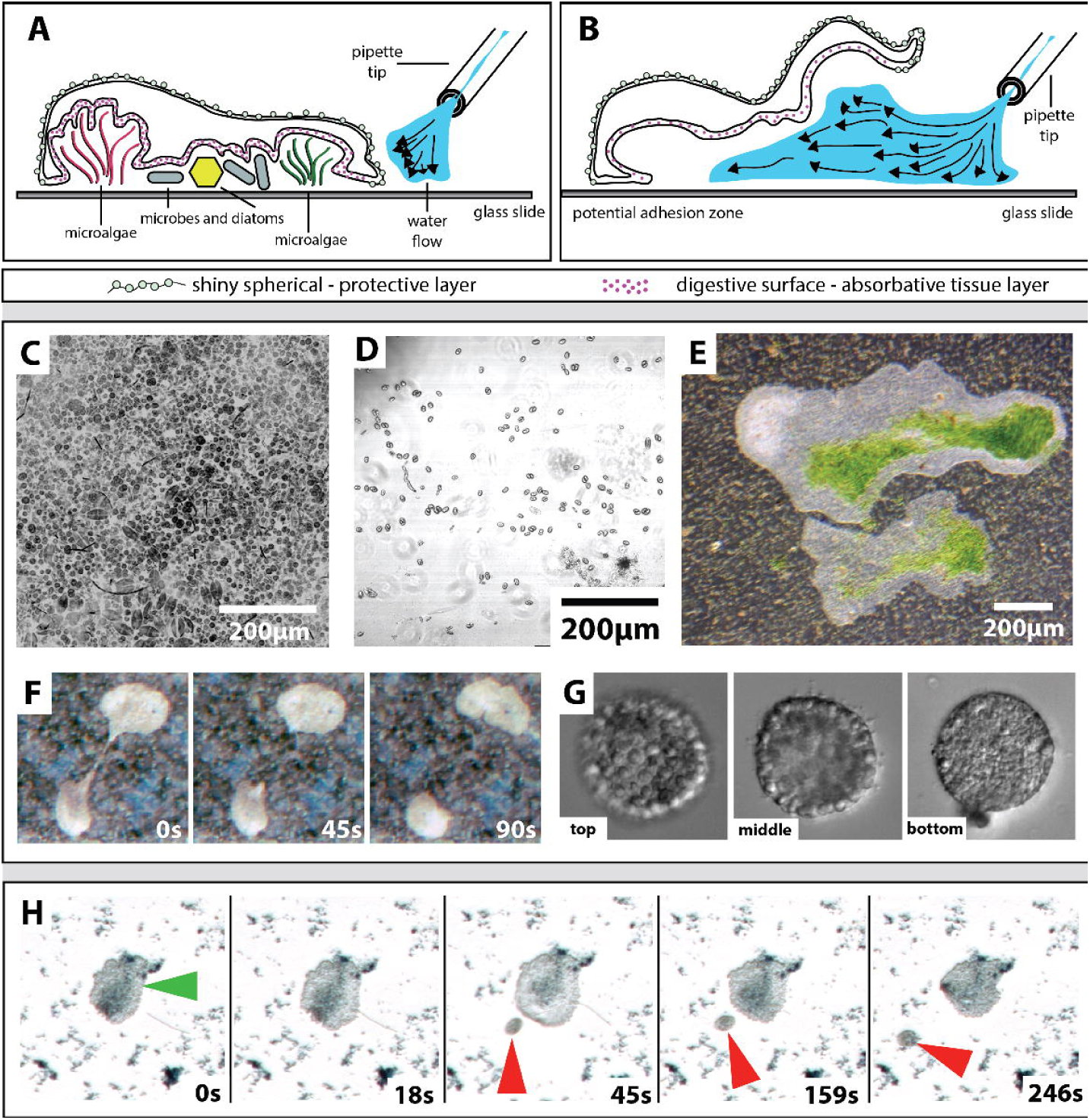
Habitat and collection of *Trichoplax adhaerens.* A) Animals adhere to the benthic environment along the thick lower epithelial layer. This layer (indicated by red dots) is thought to be involved in prey capture and digestion. The substrate in which *Trichoplax* are found is often covered with microalgae, bacteria and diatoms among other microscopic material. The upper layer of the animal consists of shiny spheres. B) Animals can be removed from their environment by applying bursts of water along the lateral edge to lift them from the surface. C) Scanning laser confocal microscopy image of a dense biofilm surface showing a diversity of unicellular organisms living among slides where placozoans grow. Each of these unicellular organisms exhibited some auto-fluorescence in the red spectrum and the image has been inverted to show contrast between each cell. D) Scattered algal cells found along the top surface of water from cultures where placozoans grow. These algal cells are imaged using DIC microscopy and also emit auto-fluorescence in the red spectrum (not shown). E) In placozoan cultures that have a high density of green algal cells, *Trichoplax* appears to “stack” cells on the top layer of themselves, although there is no evidence of feeding from this surface. F) Separation of a *Trichoplax* during asexual reproduction through binary fission, often visualized along the bottom surface of culture bowls (s=seconds). G) Three cross-sectional views of an asexually reproduced ball of cells, “swarmer”. H) A form of asexual reproduction discovered in animals living on the top surface of the water in culture. Red arrowhead indicates asexually produced clone and the green arrowhead indicates the parent.

Animals collected for this study were first collected from slides raised in sea tanks at the Kewalo Marine Laboratory (Honolulu, Hawaii). To culture animals in the lab, slides were transferred to larger containers, and over the course of one month, a dense microbial biofilm forms, consisting of “mixed” species of unicellular organisms (Figure 2c). During the weeks that follow, rapid growth of a biofilm appears in close proximity to the slide and then eventually expands to evenly coat the upper water layer along the surface tension (Figure 2d). The growth of the biofilm appears to be dependant on light availability and animals can also be cultured with minimumal light conditions (data not shown). In cultures that developed a high density of green algae, animals appear to stack cells on the top surface layer (Figure 2e) although this surface has no documented absorbative properties.

Animals found in various locations within the culture exhibited different behaviors and distinct modes of asexual reproduction. Along the bottom surface of culture bowls, animals were regularly undergoing binary fission (Figure 2f). Asexually produced “swarmers”, or animals budded from the top surface [27, 28], can regularly be found floating along the bottom surface. Swarmers have distinguishable top and bottom layers (Figure 2g), similar to adults. Animals found gliding along the top surface of the water, exhibited a unique form of asexual reproduction in which smaller clones were formed through a process of budding (Figure 2h, Videos S1-S2). Animals found along the top surface animals appear to cause a break in the surface tension (Video S1), which sometimes was associated with dispersal of an asexual clone (Video S2).

### Cellular and physiological insights into Top vs. Bottom

The top epithelial layer of the animal contains a unique cell type called shiny spheres (Figure 3a). This cell-type is thought to be involved with predator defense [15] and exhibits autofluorescence in both the blue and green spectrum (Figure 3b, Videos S3-S4). While conducting other experiments using calcium sensitive dyes, we found that exposure to high intensity near UV light or temperature shock initiated an autofluorescent wave in the FITC spectrum (Video S4). Using a temperature controller, we found that this wave of autofluorescence is typically initiated when animals are exposed to less than ~12°C or greater than 30°C (data not shown). The reaction is slowed in cold treated animals (Video S5) and expedited in hot treated animals (Video S6), although in each case animals were simulatieously exposed to close to UV light for visualization of the response. Therefore, the response could be entirely UV-related, and thermally expedited or delayed. The generation of the FITC autofluorescence appears to be in cells adjacent or beneath the shiny spheres (which lose their blue autofluorescence in the response) (Video S4). Furthermore, animals exposed to different wavelengths of light (i.e., blue, green and red), only appear to respond to light in the blue spectrum (Video S7) consistent with a previous description[29]. We hypothesize that this process is initiated by different forms of stress, but additional experiments will be required to reveal the biological significance of this peculiar behavior and molecular nature of signaling cascades involved.

**Figure 3.**
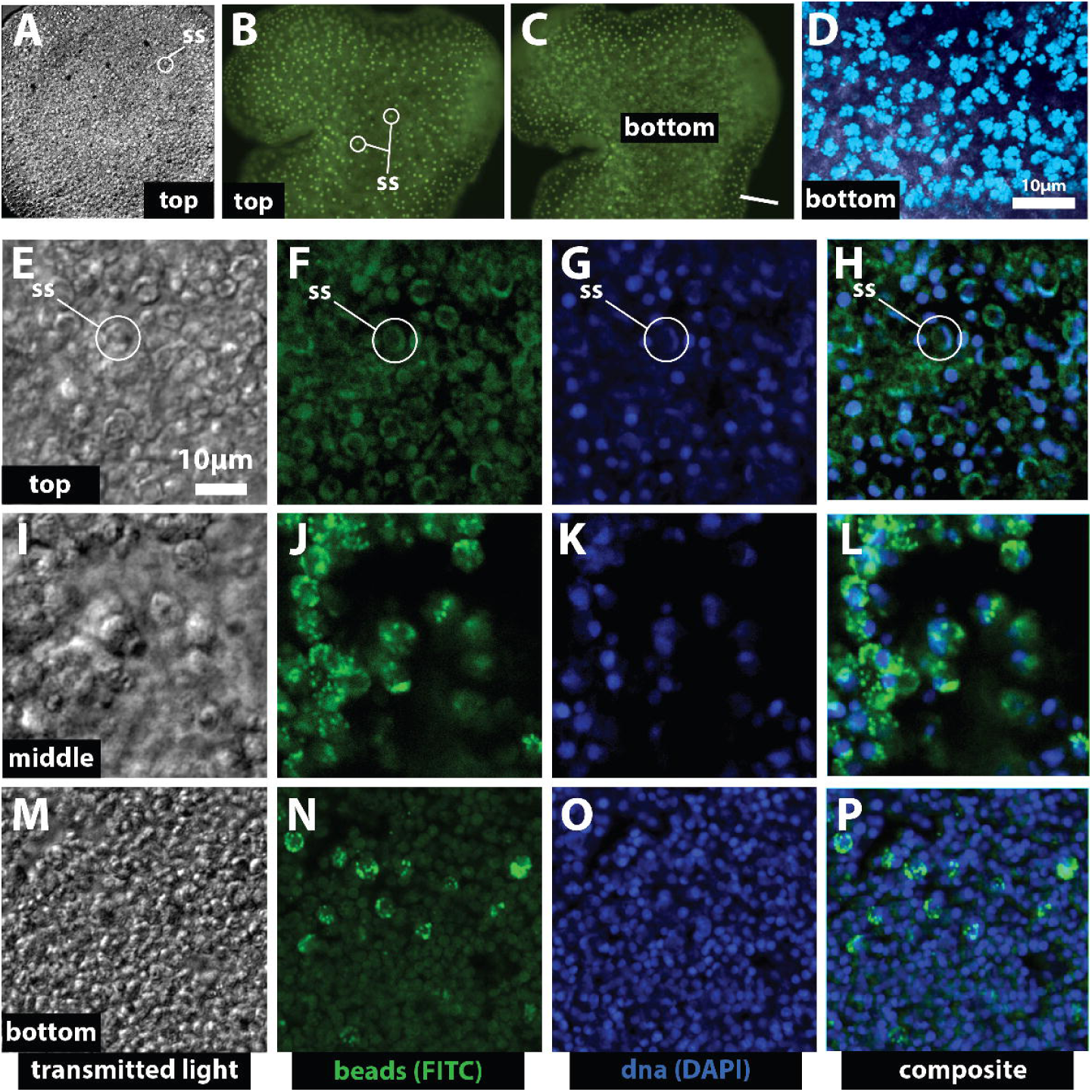
The bottom epithelial layer exhibits absorptive properties. A) The shiny spheres (SS) along the upper layer of the animal are easily visualized under transmitted light. B-C) Animal exposed to FITC wavelength of light, showing the distribution of auto-fluorescent shiny spheres along the top tissue layer (B) and absorbed beads along the bottom (C). D) Distribution of fluorescent latex beads (0.5 and 2 microns) along the lower epithelial layer after four hours of exposure. (Beads false-colored blue). E-P) Distribution of beads after twelve hours of exposure, along the upper SS region (E-H) middle fiber layer (I-L) and lower epithelial region (M-P) of an animal. Note the movement of beads from the lower layer to the middle layer from hour four (D) to hour twelve (E-P). At twelve hours, beads are found in both the lower and middle layers. Nuclei are shown for scale.

Placozoans are thought to feed on bacteria, yeast, algae or some by-product of biofilms through absorbtion along the bottom epithelial layer[11, 26, 30–32]. To test the absorbative properties of the bottom layer, autofluorescent latex beads of 0.5 and 2 microns were incubated with natural biofilms to create a bead/biofilm matrix. We exposed animals to these beads to see if they would be absorbed by *Trichoplax.* At low magnification, the autofluorescence of the shiny speres was observed at the FITC wavelength (Figure 3b). As we adjusted the focal plane to focus on underlying cells (Figure 3c), we observed the fluorescence from absorbed beads (Video S3). Under higher magnifications, we observed shiny spheres along the top surface (Figure 3e), and were able to differentiate two different diameter beads that were compartmentalized within cells along the bottom layer of the animal (Figure 3f, false-colored blue). Additionally, we observed foraging Trichoplax clearing a biofilm using longterm timelapse microscopy (Video S8). These findings suggest that the bottom layer has both digestive[26] and absorbative properties.

### Genomic aspects of placozoans

Since their first discovery, placozoans have been described as diploblastic animals that exhibit a dorso-ventral polarity along the top and bottom tissue layers. To test if this axis is driven by the same set of transcription factors that define doral-ventral patterning in bilaterians [33–37], we characterized the Chordin TgfB signaling pathway in *Trichoplax.* One of the primary antagonistic iterplays in this pathway is between Bmp ligands and Chordin genes[33–35, 37]. Genomic evidence had suggested that a chordin-like gene is present in the *Trichoplax* genome[3, 13], yet several elements of the TgfB pathway have yet to be phylogenetically resolved [38–42]. We conducted phylogenetic analyses using sequences, domains, and genomic synteny to better classify these genes.

Vertebrate chordin genes consist of multiple cysteine-rich (CR) repeats linked to a series chordin (Chd) domains. Functional experiments have shown that the CR repeats of vertebrate Chordin (primarily CR1 and CR3) appear to confer the dorsalizing activity of the gene during development [34]. There are a number of other genes containing CR domain repeats[43] and some of these also exhibit asymmetric patterning activity similar to chordin in other animals[34, 44]. Our phylogenetic analyses of CR repeats from a diverse set of species, suggests that *Trichoplax* has a Chordin-like gene that lacks a recognizeable Chd domain (Figure 4a-b). The predicted Chd domains in each Chordin protein appear to be quite variable across the animal kingdom (Figure S1). In addition, the relationship between bona fide chordin genes and some non-bilaterian CR-domain containing genes remain unresolved (e.g. ctenophores [40], sponges [45], and *Hydra* [46]) (Figure 4a). We used synteny, or the comparison of gene location between the genomes of two species (in this case *Trichoplax* and the cnidarian, *Nematostella vectensis*), as evidence for orthology of *Chordin* from *Trichoplax.* This approach has been applied in many other studies to help resolve gene orthology [3, 47–50]. The scaffolds containing *Chordin* in *Trichoplax* and *Nematostella* share four genes with each other, flanking both sides of the gene in each animal (Figure 4c). Together, this evidence suggests that *Trichoplax* has a Chordin gene that lacks a recognizeable Chd domain but contains 4 CR repeats.

**Figure 4.**
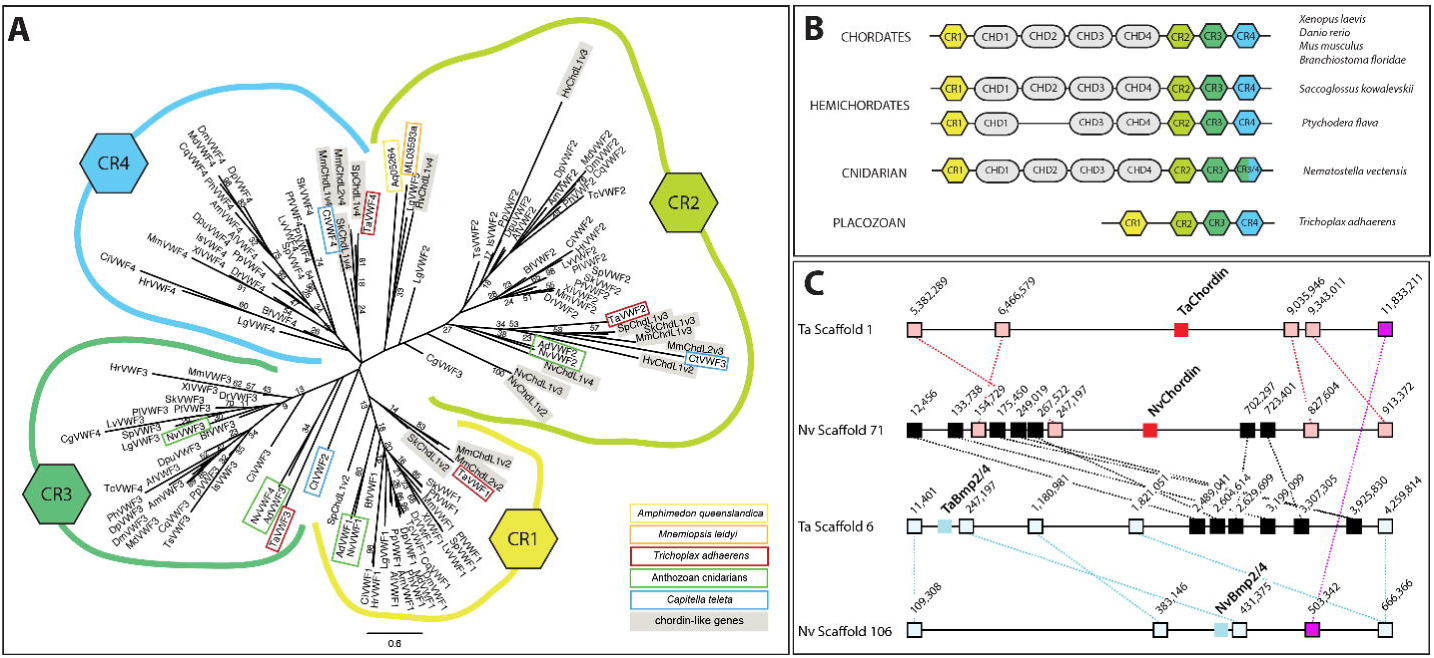
*Trichoplax* has a chordin gene consisting of only CR domains. A) Maximum likelihood analysis of diverse CR domains from Chordin and Chordin-related proteins. *Trichoplax* CR domains group with the four different CR domains of other Chordin genes. B) Schematic diagram of Chordin (Chd) and cysteine rich (CR) domains found among animals. C) Synteny analysis based on at least 1000 base-pair regions of conservation between scaffolds of *Trichoplax* and *Nematostella.* Scaffolds of Chordin and Bmp2/4 that share conserved regions are identified by dotted lines. Vertical numbers indicate the position along the scaffold.

Chordin is known to functionally interact with TgfB signaling to specificy the dorsal-ventral axis of diverse bilaterians. Phylogenetic analysis of the TgfB complement of *Trichoplax* is less clear. A number of conditions were tested (see methods) yet no tree clearly resolved all *Trichoplax* Bmp ligands (Figure S2). Again, synteny analysis between the cnidarian *Nematostella vectensis* and *Trichoplax* confirms the existence of linkage between orthologs of *Bmp2/4, Gdf5* and *Bmp5/8* (Figure 4c, Figure S3). We identified a *Trichoplax* gene related to *BMP3* that is also present in ctenophores[40] and may be a distant relative to both *BMP3* and *Admp* (and potentially *Nodal*), however, we were unable to confirm the orthology of this gene with synteny.

### Elaborate gene expression patterns of key developmental transcription factors

We analyzed a diverse set of eight different genes to determine if territorial patterning occurs and if dorsal-ventral patterning genes are expressed along the top-bottom axis of *Trichoplax. Beta-actin,* a house-keeping gene which often exhibits ubiquitous expression in other species, is expressed throughout the *Trichoplax* body in both top and bottom tissue layers (Figure 5A). *Trichoplax* has a single ortholog of the zinc-finger *Snail* transcription factor[51] (a marker for mesoderm in bilaterians[52, 53] and endomesodermal cells in cnidarians) [54–56]. *Snail* is broadly expressed in the bottom tissue layer and exhibits a salt-n-pepper pattern along the top surface (Figure 5B). *BMP2/4* is expressed primarily along the bottom tissue layer, with a few scattered cells around the lateral edge (Figure 5C). *Chordin,* a Bmp antagonist, is found along the bottom surface (Figure 5D) and localizes to an overlapping region as *BMP2/4* (Figure 5E). *Bmp3* is expressed asymmetrically along one lateral edge of the animal (Figure 5F). *Gdf5* is expressed in a ring around the bottom layer (Figure 5G). A single *Elav* gene, a broad-neuronal marker in *Nematostella* [57, 58], is found expressed throughout the lower epithelial layer (Figure 5H). The ParaHox gene *GSX* (also called *Trox-2* [16]) is broadly expressed throughout the lower epithelial layer (Figure 5I).

**Figure 5.**
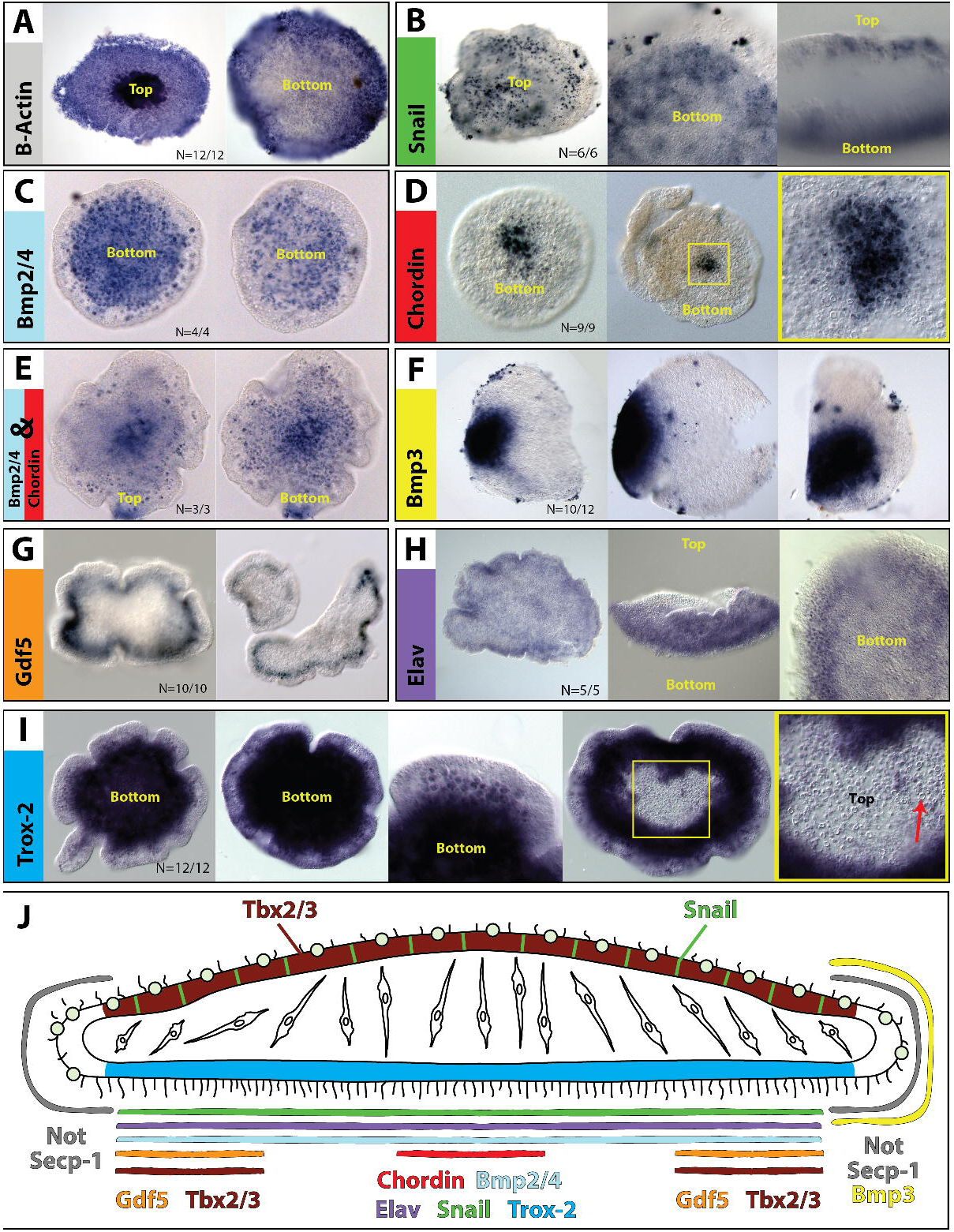
Asymmetric expression of mRNA transcripts along the body of *Trichoplax.* A) *Beta-actin* is expressed throughout the tissue layers and appears highly expressed or in a greater number of cells when the animal is contracted (top, left) B) *Snail* is expressed in a punctate (salt-and-pepper) pattern along the upper/lower layers C) *Bmp2/4* is expressed in the bottom tissue layer. D) *Chordin* is expressed in a small subset of cells along the bottom layer. E) *Chordin* and *Bmp2/4* are in overlapping domains along the bottom tissue layer. F) Bmp3 is expressed along one side of the animal. G) *Gdf5* is expressed in a ring around the bottom layer. H) *Elav* is found broadly along the lower layer. I) *Trox-2* is highly expressed throughout the bottom layer. I) Summary of expression domains of genes from within this manuscript and previous expressions of *Tbx2/3*[21], *Secp-1* and *Not*[22].

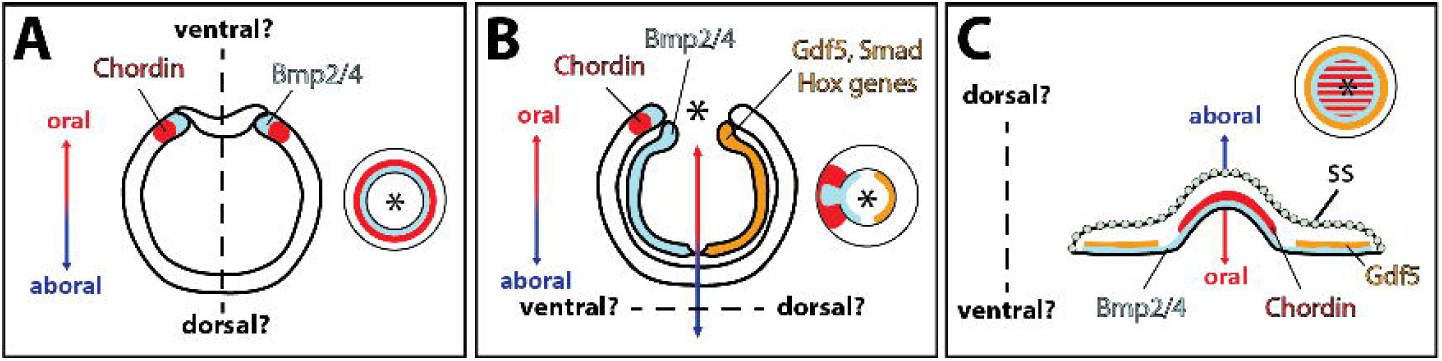

## Discussion

Much has been learned about Placozoan biology through the application of electron microscopy techniques [11, 17, 27, 30, 59–64]. On the other hand, to date much less has been gleaned through gene expression studies despite the first gene expression pattern being reported fourteen years ago [21] and the genome being published nine years ago[3]. In this study we establish a new *in situ* hybridization protocol and use this protocol to show elaborate expression patterns for a range of important developmental patterning genes. These data show a level of complexity in conflict with the low number of morphologically distinct cell-types (approximately six) reported in *Trichoplax* [3, 12, 19].

The body axis of *Trichoplax* is overtly similar to the oral-aboral axis of cnidarians. In the anthozoan cnidarian *Nematostella,* the transcription factor *Snail* is expressed throughout the endomesoderm of the animal [54, 56, 65], and the Chordin-TgfB pathway patterns the directive axis, perpendicular to the oral-aboral axis of the animal [66–69]. Although the Chordin-TgfB patterning is perpendicular, the signaling begins along the animal pole of the embryo, which is the future oral side of the animal. The earliest expression domain of both *Chordin* and *Bmp2/4* is radial along the site of gastrulation[38, 70–72] (Figure 6a). During gastrulation, the expression domains become asymmetric and are opposite of *Gdf5, Smad1/5,* and several *Hox* genes among other factors [38, 66–69, 71, 73, 74] (Figure 6b). The change from radial to asymmetric patterning in *Nematostella* begs the question of whether the dorsal-ventral axis is parrallel to the oral-aboral axis (Figure 6a) or perpendicular (Figure 6b). Across cnidarians, *Bmp2/4* appears asymmetric around the blastopore and into larval development [75, 76], but it is currently unclear if these expression domains start out as exhibiting radial symmetry.

This study shows that orthologs of *Chordin, Bmp2/4, Gdf5, Elav* and Gsx (*Trox-2) Trichoplax* are all expressed along bottom layer in at least three territories (Figure 5J). In addition, the homeobox T-box transcription factor *Tbx2/3* has previously been shown to be expressed in the top and bottom layer[21]. In the ctenophore *Mnemiopsis leidyi, Tbx2/3* is primarily expressed in the aboral region[77] with faint expression along the oral surface. Likewise, Tbx2/3 is primarily expressed along the oral surface of the cnidarian *Nematostella vectensis* (Figure S4). These expression data along with the diverse studies on the feeding behaviour of *Trichoplax,* suggest that the bottom surface of *Trichoplax* is homologous to the endomesoderm of cnidarians, and thus the “dorso-ventral” axis of *Trichoplax* is homologous to the oral-aboral axis of cnidarians.

The most detailed analyses of the emergence of dorsal-ventral patterning has focused on the cnidarian *Nematostella vectensis.* Expression data suggest that this pathway patterns the directive axis of *N. vectensis,* but it is unclear whether this axis is homologous to the bilaterian dorsal-ventral axis [42]. Comparing the totality of evidence to date, one could argue that the common ancestor of placozoans and cnidarians (i.e., the last common parahoxazoan [78]), could have exhibited: a) radial symmetry of *Chordin* and *Bmp2/4* similar to placozoans, or b) asymmetric expression of the two genes. In the latter scenario, placozoans would had to have lost components of the cnidarian directive axis. Due to the number of similarities between the bottom surface and oral or endo/mesoderm patterning genes of cnidarians (e.g. expression of *Chordin, Bmp2/4, Gdf5, Elav, Gsx* and broad expression of *Snail*) it is our interpretation that the common ancestor was a radial animal. Furthermore, we propose that these expression patterns of Trichoplax are more comparable to blastula patterning in Bilateria, and that the directive axis patterning in Nematostella vectensis was derived within Anthozoa.. This would further imply that the asymmetric patterning of Chordin/Bmp in cnidarians and bilaterians was independantly evolved. Unfortunately, until it becomes possible to observe embryogenesis in *Trichoplax,* it will be impossible to test the role of these genes during embryonic development.

Our results suggest that the *Gsx* ortholog, *Trox-2,* is expressed throughout the bottom layer of the animal, rather than in a ring as previously reported [16]. In cnidarians [79–81], a sea urchin [82], two mollusks [83], and numerous chordates [84–87], *Gsx* orthologs have been implicated in patterning neurons. Previous studies have suggested that the lower layer of *Trichoplax* is a simplified digestive surface [26, 30–32]. Interestingly, the expression of neuronal patterning genes in digestive tissue is reminiscent of patterns seen in Cnidaria. The larval and adult nervous system of anthozoan cnidarians is both ectodermal and endomesodermally derived. A large population of endomesodermal derived neurons are comprised of sensory-like morphology, express *Elav1* and follow along parietal muscles of the digestive track[57, 58, 88]. In Hydrozoan cnidarians, the nervous system is thought to be derived from I-cells that orginate in the endomesoderm (gastroderm) during embryonic development [89–91], although sensory cells can emerge from I-cell free animals [92, 93]. In the absence of a true nervous (or musculature) system, neuropeptide and calcium signaling play a role in complex behaviors of these animals [94, 95]. The presence of this diverse set of markers and digestive cells along the bottom surface provide additional support for the bottom surface of *Trichoplax* being related to cnidarian oral and endomesoderm tissue. Furthermore the bottom/oral side of the animal is a multifunctional surface, capable of particle transport (e.g. latex beads) and potentially sensory function (e.g. expression of *Trox-2* and *Elav*).

Our findings, although suggestive, require greater analysis and further study to truly understand how these and other developmental genes shape the body of *Trichoplax.* For example, the fiber layer of these animals has been overlooked in all studies to date and may require better sampling techniques (e.g. histological sectioning) to completely identify transcripts localized to this region. This research has provided a crucial protocol for advancing our knowledge of these egnimatic understudied animals. Few identified cells types, yet a diverse molecular toolkit make us ask the question, “where have they hidden their morphological complexity?” One possibility is that the “placula/swarmer” morphotypes, currently cultured in labs around the world, are simply the larval or dispersal stage of this clade and are unable to produce viable gametes. Efforts should be made to identify the complete life cycle of these organisms. For example, the adult phase could be a cryptic or parasitic form that bears no morphological similarities to the larval form. After all, an entire clade of lophotrochozoan bilaterians, the Dicyemids, are internal parasites of cephalopod nephridia[96, 97]. It would not be a surpise to learn that *Trichoplax* sequences show up in some future metagenic sequencing projects that were not even aimed at finding cryptic species. Hopefully, we will soon know the answer to this “simple” problem.

## Methods

### Collection/culture of *Trichoplax*

All animals were collected from an outdoor flow-through sea water system at the Kewalo Marine Laboratory (Honolulu, HI). Seawater tanks were seeded with heavily biofouled rocks and algae samples from the nearby Kewalo basin. Animals were collected from glass slides that were placed in a metal slide rack and submerged with the rocks/algae for 3-5 weeks. Tanks were kept under constant low flow rates in full sunlight. Animals were typically found creeping along slides covered with a dense bacterial biofilm that also had microalgae and diatoms present (Figure 2a/c, Video S8). Specimen were removed from the surface of the slide by gentle pipette pressure (Figure 2b) and transferred to gelatin-coated plastic dishes. Gelatin coated dishes were prepared by disolving gelatin (1x) into distilled water and covering each petri dish (47mm) with approximately 2 ml of liquid. Excess gelatin was then removed and dishes were dried overnight. After transfer, animals were kept for twenty-four hours to allow for acclimation to the dish.

### Latex bead experiments

Animals were kept for longer periods of time (e.g. bead feeding experiments) in glass bowls and provided a drop of Micro Algae Grow (http://florida-aqua-farms.com) every 2-3 days. We used latex beads of 0.5 and 2 microns in diameter (Cat.# L3280 & L4530, Sigma Aldrich) to cover the surface of plastic petri dishes filled with seawater. Beads were suspended at a dilution of 1:100 from its stock in seawater, then two droplets from both bead samples were added to each dish and left to settle on the bottom overnight. The next day, dishes were washed with seawater to remove excess floating beads. Animals were then added to each dish and allowed to come in contact with the beads along the surface of the dish for four hours. After four or twelve hours of exposure to beads, animals were fixed and moved to glass slides for visualization by microscopy.

### Imaging

All experiments, except those involving fluorescent beads, were visualized using an Axioscope 2 compound microscope with an AxioCam (HRc) camera. Images were compiled using Axiovision software (Zeiss Inc, Jena, Germany). Autofluorescence of shiny sphericals could be seen using a FITC filter cube and was visualized using 488 wavelength settings. For fluorescent bead experiments, live animals were placed on glass slides and visualized using a Zeiss 710 scanning laser confocal. Images were cropped and assembled using Adobe Photoshop and Illustrator CS6.

### Autofluorescent dynamics experiments

Animals were imaged using a newly developed preparation. In brief, a single animal was mounted on a plastic or glass-bottom 35mm Petri dish in ~50mkl high viscosity solution, covered with glass coverslip, sealed with Vaseline and placed on the stage of an inverted microscope. Such “microchamber” conditions substantially restricted both lateral and vertical movements allowing prolong imaging of live animals. High viscosity solution contained: filtered sea water; methylcellulose X-Ymg per ml.

Water was applied using gravity fed perfusion system. Temperature of the water was controlled by bipolar temperature controller (model CL-100, Warner Instruments) and a SC-20 dual in-line solution heater-cooler (Warner Instruments, Harvard Apparatus). Temperatures in Petri dish were measured using a TA-29 thermistor (Warner instruments). Both perfusion output ports and the external thermistor were positioned in the close proximity to the imaging area.

Fluorescence imaging was performed on an inverted microscope (Olympus IX-71) equipped with a cooled CCD camera (ORCA R2, Hamamatsu) under the control of Imaging Workbench 6 software (INDEC Systems). A standard FITC or Fura-2 filter set (excitation at 510 nm, emission at 530 nm and excitation at 340 nm or 380 nm, emission at 510 nm) were used for measurements of endogenous fluorescence intensity. Images were collected at the rate ~1, 2 or 4Hz (specified individually for each data set). Recorded data were stored as image stacks, analyzed off-line using Imaging Workbench 6 or ImageJ 1.42 (available from public domain at http://rsbweb.nih.gov/ij/index.html).

### RNA, cDNA and Gene Isolation

We collected a total of approximately 100 animals from biofilm slides and transferred animals to gelatin coated dishes. Any debris transferred with animals was removed periodically to eliminate foreign contaminants from our samples. Animals were starved overnight and cleaned again prior to RNA collection. All animals were transferred to a 1.5 ml tube and gently centrifuged (3000g) to the bottom of the tube. All remaining seawater was removed and animals were suspended in Trizol (Cat.# 15596-026, Invitrogen) and vortexed to lyse the tissue. Total RNA was extracted using the manufacturers protocol. Samples were DNAse (Cat.# 79254, Quiagen) treated to remove DNA contamination (15 minutes at 37°C) and then an additional phenol choloroform extraction was used to clean and precipitate the RNA. cDNA was constructed using a SMARTer^®^ RACE cDNA Amplification Kit (Cat.# 634923, ClonTech) for RACE cDNA and Advantage^®^ 2 PCR Kit (Cat.#639206, ClonTech) for cDNA to be used for general PCR.

A number of desirable genes for analysis were identified using the genome [3] through the Joint Genomic Institute website (http://genome.jgi-psf.org/Triad1/Triad1.home.html). Nucleotide sequences were gathered and primers designed using MacVector (www.macvector.com). A list of primers used in this study can be found in Table S1. Fragments of approximately 600-1100 bases were cloned using the pGEM^®^ T Easy System (Cat.# A1360, Promega). Probes were synthesized using the MEGAscript^®^ T7/SP6 Transcription Kits (Cat.# AM1334, AM1330, Invitrogen) and were stored at −20°C.

### Phylogenetic and synteny analysis

Protein sequences from a diverse set of taxa were collected using the NCBI protein databases (http://www.ncbi.nlm.nih.gov). A list of the different species used in our phylogenetic analyses can be found in Figure S2. Genomic resources of *Mnemiopsis leydei* [98], *Acropora digitifera* [99], *Capitella teleta, Lottia gigantica, Helobdella robusta* [50], *Daphnia pulex* [100] and *Branchiostoma floridae* [101] were used in this study. Protein coding domains were predicted using SMART [102] and all sequences were aligned using MUSCLE [103]. Trees were constructed using MrBayes [104, 105] using five independent runs, consisting of 5,000,000 generations using “mixed” models. A second tree was constructed using maximum-likelihood analysis using RaxML (version 7.2.8) as described [98]. Maximum-likelihood tree bootstraps were based on 100 replicates. Four different allignments were run using each analsysis. They included conditions listed below each tree. Trees were imported and edited using FigTree (version 1.4.0) [106].

### Whole mount *in situ* hybridization

Animals were transferred from glass slides to gelatin-coated dishes using a glass pasteur pipette (Cat.#CLS7095D5X, Sigma-Aldrich) and allowed to settle on the bottom of the dish overnight. Fixation was achieved by gently adding ice-cold fix (4% PFA, 0.2% Glutaraldehyde in high salt seawater; 0.5g NaCl/50mL seawater) to the dish for 90 seconds. Following this, the solution was gently removed and ice-cold 4% PFA (in high salt seawater) was added and dishes placed at 4°C for 1 hour. Fix was removed and animals were washed three times in ice-cold DEPC-treated H_2_O, then dehydrated to 100% methanol over several steps (25% methanol:DEPC-H_2_O for 5 minutes, 50% methanol:DEPC-H_2_O for 5 minutes and 75% methanol:DEPC-H_2_O for 5 minutes). Animals were washed in 100% methanol for one hour at 4°C and then used immediately for *in situ* hybridization.

Animals were transferred to a 96-well plate (1 animal per well), and rehydrated into 1x PBS pH7.4 + 0.1% Tween-20 (PTw) using a series of washes (150 μl per wash; 75% methanol:25% PTw, 50% methanol:50% PTw, 25% methanol:75% PTw), before washing 5 times in PTw. Proteinase K was added (0.01mg/mL) for 5 minutes before digestion was stopped by two, 5-minute washes in PTw+2mg/mL glycine. Animals were then washed twice (5 minutes each time) in 0.1% triethanolamine:PTw This solution was substituted with 0.1% triethanolamine:PTw with 3μL/mL acetic anhydride added for 5 minutes, followed by a 5 minute wash in 0.1% triethanolamine:PTw with 6μL/mL acetic anhydride added. Animals were then washed 2 times for 5 minutes each in PTw and refixed in 4% PFA in PTw at 4°C for 1 hour. Fix was removed and animals washed 5 times in PTw, and then 2 times (10 minutes each wash) in hybridization buffer (50% deionized formamide, 5x SSC pH 4.5, 50g/mL heparin, 0.5% Tween-20 freshly made, 1% SDS, 100g/mL salmon sperm DNA). Animals were prehybridized in hybridization buffer for 3 hours to overnight at hybridization temperature. One crucial difference in our protocol, compared to previously published data, is that we hybridized our probes at 64°C, where all other published works used 50°C. Digoxygenin (DIG)-labelled anti-sense RNA probe was added to a final concentration of 1ng/L and left to incubate at hybridization temperature for 48 hours. Following hybridization, probe was removed and a series of washes conducted at hybridization temperature; 10 minutes in 100% hybridization buffer, 20 minutes in 100% hybridization buffer, 20 minutes in 75% hybridization buffer:25% 2x SSC, 20 minutes in 50% hybridization buffer:50% 2x SSC, 20 minutes in 25% hybridization buffer:75% 2x SSC, 3 × 20 minute washes in 2x SSC and 3 × 10 minute washes in 0.05x SSC. A series of wash steps at room temperature were then conducted; 5 minutes in 75% 0.05x SSC:25% PTw, 5 minutes in 50% 0.05x SSC:50% PTw, 5 minutes in 25% 0.05x SSC:75% PTw and 3 × 10 minute washes in PTw. Samples were incubated in 1x Roche Blocking reagent:maleic acid buffer (Cat.# <u>11096176001</u>) overnight (at 4°C?), then incubated in anti-DIG-AP antibody (catalog number 11093274910, 1:5000) overnight at 4°C. The following day, tissue was washed in at least 10 × 30 minute washes in PTw at room temperature, followed by 3 × 5 minute washes in PBS. Animals were then washed for 2 × 5 minute washes in AP-M buffer (0.1M NaCl, 0.1M Tris pH9.5, 0.05% Tween-20), then 2 × 5 minute washes in AP buffer (0.1M NaCl, 0.1M Tris pH9.5, 0.05% Tween-20, 0.1M MgCl_2_). Visualization of probe was achieved by adding 3.3μL/mL NBT (US biological N2585; 75mg/mL in 70% dimethylformamide) and 3.3μL BCIP (US biological B0800; 50mg/mL in dimethylformamide) to AP buffer and incubating at room temperature until the desired level of staining intensity was reached. The reaction was stopped by washing in PBS or PTw for 3-5 times, and animals mounted in 70% glycerol:PBS for analysis.

## Authors’ contributions

TQD collected and reared animals, conducted experiments. TQD and JF ran all phylogenetic analysis. TQD and MQM were involved in project design. TQD, JR and MQM were involved in synthesis of the final manuscript.

## Acknowledgements

TQD would like to thank Michael G. Hadfield for introduction and encouragement to work with the animal, *Trichoplax.* We would also like to thank Vicky Pearse for her insight and expertise. This research was funded by research grants obtained by MQM from both NIH and NSF.

## Supplemental Resources Figure Legends

**Figure S1** – Chordin genes across the animal kingdom. Diversity of cysteine-rich (CR) and chordin (Chd) domains among chordin and chordin-related genes found within animals. Schematic created based on the phylogenetic-relatedness of identified domains with these animal groups (Figure S2). Chordin-related genes contain additional domains (e.g. IBM, Kazal, Till, KU) and their CR domains are all more closely related to CR2 rather than other CR domains.

**Figure S2** – Species names and files associated with all phylogenetic analysis within the manuscript

**Figure S3** – Synteny analysis showing regions of conservation between *Nematostella* and *Trichoplax* TgfB related genes.

**Figure S4** – Tbx2/3 expression during early development of *Nematostella vectensis*. (* Indicates the oral pole)

**Video S1** – Animals on the water surface appear to cause a break in surface tension, caused by unknown reason.

**Video S2** – Animal along the water surface undergoing asexual reproduction

**Video S3** – Video showing different focal planes of an animal exhibiting auto-fluorescent shiny sphericals and fluorescence of latex beads taken up on the oral surface

**Video S4** – Auto-fluorescent induced phenomenon of shiny spherical cells

**Video S5** – Video showing auto-fluorescent response of shiny spherical cells to progressively colder temperature

**Video S6** – Video showing auto-fluorescent response of shiny spherical cells to progressively hotter temperature

**Video S7** – Behavior response to different wavelengths of light

**Video S8** – Time-lapse of asymmetric feeding behavior

## References

1. Pearse VB, Voigt O: Field biology of placozoans (Trichoplax): distribution, diversity, biotic interactions. Integr Comp Biol 2007, 47:677–92.

2. Eitel M, Osigus H-JJ, DeSalle R, Schierwater B: Global diversity of the Placozoa. PLoS One 2013, 8:e57131.

3. Srivastava M, Begovic E, Chapman J, Putnam NH, Hellsten U, Kawashima T, Kuo A, Mitros T, Salamov A, Carpenter ML, Signorovitch AY, Moreno M a, Kamm K, Grimwood J, Schmutz J, Shapiro H, Grigoriev I V, Buss LW, Schierwater B, Dellaporta SL, Rokhsar DS: The Trichoplax genome and the nature of placozoans. Nature 2008, 454:955–60.

4. Hejnol A, Obst M, Stamatakis A, Ott M, Rouse GW, Edgecombe GD, Martinez P, Baguñà J, Bailly X, Jondelius U, Wiens M, Müller WEG, Seaver E, Wheeler WC, Martindale MQ, Giribet G, Dunn CW: Assessing the root of bilaterian animals with scalable phylogenomic methods. Proc Biol Sci 2009, 276:4261–4270.

5. Philippe H, Derelle R, Lopez P, Pick K, Borchiellini C, Boury-Esnault N, Vacelet J, Renard E, Houliston E, Quéinnec E, Da Silva C, Wincker P, Le Guyader H, Leys S, Jackson DJ, Schreiber F, Erpenbeck D, Morgenstern B, Wörheide G, Manuel M: Phylogenomics revives traditional views on deep animal relationships. Curr Biol 2009, 19:706–12.

6. Borowiec ML, Lee EK, Chiu JC, Plachetzki DC: Dissecting phylogenetic signal and accounting for bias in whole-genome data sets supports the Ctenophora as sister to remaining Metazoa. BMC Genomics 2015, 16:1–15.

7. Simion P, Philippe H, Baurain D, Jager M, Richter DJ, Di Franco A, Roure B, Satoh N, Quéinnec É, Ereskovsky A, Lapébie P, Corre E, Delsuc F, King N, Wörheide G, Manuel M: A Large and Consistent Phylogenomic Dataset Supports Sponges as the Sister Group to All Other Animals. Curr Biol 2017, 27:1–10.

8. Dellaporta SL, Xu A, Sagasser S, Jakob W, Moreno M a, Buss LW, Schierwater B: Mitochondrial genome of Trichoplax adhaerens supports placozoa as the basal lower metazoan phylum. Proc Natl Acad Sci U S A 2006, 103:8751–6.

9. Schierwater B, Eitel M, Jakob W, Osigus HJ, Hadrys H, Dellaporta SL, Kolokotronis SO, DeSalle R: Concatenated analysis sheds light on early metazoan evolution and fuels a modern “urmetazoon” hypothesis. PLoS Biol 2009, 7.

10. Grell K: Trichoplax adhaerens F. E. Schulze und die Entstehung der Metazoen. Naturw Rundschau 1971, 24:160–161.

11. Grell, KG, Benwitz G: Die Ultrastruktur von Trichoplax adhaerens F.E. Schulze. Cytobiology 1971, 4:216–240.

12. Grell K: Eibildung und Furchung von Trichoplax adhaerens FE Schulze (Placozoa). Zoomorphology 1972, 73:297–314.

13. Richards GS, Degnan BM: The dawn of developmental signaling in the metazoa. Cold Spring Harb Symp Quant Biol 2009, 74:81–90.

14. Ringrose JH, van den Toorn HWP, Eitel M, Post H, Neerincx P, Schierwater B, Altelaar a FM, Heck AJR: Deep proteome profiling of Trichoplax adhaerens reveals remarkable features at the origin of metazoan multicellularity. Nat Commun 2013, 4(May 2012):1408.

15. Jackson AM, Buss LW: Shiny spheres of placozoans (Trichoplax) function in anti-predator defense. Invertebr Biol 2009, 128:205–212.

16. Jakob W, Sagasser S, Dellaporta S, Holland P, Kuhn K, Schierwater B: The Trox-2 Hox/ParaHox gene of Trichoplax (Placozoa) marks an epithelial boundary. Dev Genes Evol 2004, 214:170–5.

17. Guidi L, Eitel M, Cesarini E, Schierwater B, Balsamo M: Ultrastructural analyses support different morphological lineages in the phylum Placozoa Grell, 1971. J Morphol 2011, 272:371–8.

18. Pearse VB, Uehara T, Miller RL: Birefringent Granules in placozoans (Trichoplax adhaerens). Trans Am Microsc Soc 1994, 113:385–389.

19. Smith CL, Varoqueaux F, Kittelmann M, Azzam RN, Cooper B, Winters CA, Eitel M, Fasshauer D, Reese TS: Novel Cell Types, Neurosecretory Cells, and Body Plan of the Early-Diverging Metazoan Trichoplax adhaerens. Curr Biol 2014, 24:1–8.

20. Schuchert P: Trichoplax adhaerens (Phylum Placozoa) has cells that react with antibodies against the neuropeptide RFamine. Acta Zool 1993, 74:115–117.

21. Martinelli C, Spring J: Distinct expression patterns of the two T-box homologues Brachyury and Tbx2/3 in the placozoan Trichoplax adhaerens. Dev Genes Evol 2003, 213:492–9.

22. Martinelli C, Spring J: Expression pattern of the homeobox gene Not in the basal metazoan Trichoplax adhaerens. Gene Expr Patterns 2004, 4:443–7.

23. Hadrys T, DeSalle R, Sagasser S, Fischer N, Schierwater B: The Trichoplax PaxB gene: a putative Proto-PaxA/B/C gene predating the origin of nerve and sensory cells. Mol Biol Evol 2005, 22:1569–78.

24. Signorovitch AY, Dellaporta SL, Buss LW: Caribbean placozoan phylogeography. Biol Bull 2006, 211:149–56.

25. Eitel M, Schierwater B: The phylogeography of the Placozoa suggests a taxon-rich phylum in tropical and subtropical waters. Mol Ecol 2010, 19:2315–27.

26. Smith CL, Pivovarova N, Reese TS: Coordinated feeding behavior in trichoplax, an animal without synapses. PLoS One 2015, 10:1–15.

27. Thiemann M, Ruthmann A: Trichoplax adhaerens Schulze, F. E. (Placozoa) - The formation of swarmers. Z Naturforsch C 1988, 43:955–957.

28. Thiemann M, Ruthmann A: Alternative modes of asexual reproduction inTrichoplax adhaerens (Placozoa). Zoomorphology 1991:165–174.

29. Pearse VB: Growth and Behavior of Trichoplax adhaerens: First Record of the Phylum Placozoa in Hawaii. Pacific Sci 1989, 43:117–121.

30. Wenderoth H: Transepithelial cytophagy by trichoplax adhaerens placozoa feeding on yeast saccharomyces cerevisiae. Z Naturforsch 1986, 41C:343–347.

31. Ueda T, Koya S, Maruyama YK: Dynamic patterns in the locomotion and feeding behaviors by the placozoan Trichoplax adhaerence. Biosystems 1999, 54:65–70.

32. Smith CL, Reese TS: Adherens Junctions Modulate Diffusion between Epithelial Cells in Trichoplax adhaerens. Biol Bull 2016, 231:216–224.

33. Sasai Y, Lu B, Steinbeisser H, Geissert D, Gont LK, De Robertis EM: Xenopus chordin: a novel dorsalizing factor activated by organizer-specific homeobox genes. Cell 1994, 79:779–90.

34. Larraín J, Bachiller D, Lu B, Agius E, Piccolo S, De Robertis EM: BMP-binding modules in chordin: a model for signalling regulation in the extracellular space. Development 2000, 127:821–30.

35. van der Zee M, Stockhammer O, von Levetzow C, Nunes da Fonseca R, Roth S: Sog/Chordin is required for ventral-to-dorsal Dpp/BMP transport and head formation in a short germ insect. Proc Natl Acad Sci U S A 2006, 103:16307–12.

36. Lowe CJ, Terasaki M, Wu M, Freeman RM, Runft L, Kwan K, Haigo S, Aronowicz J, Lander E, Gruber C, Smith M, Kirschner M, Gerhart J: Dorsoventral patterning in hemichordates: insights into early chordate evolution. PLoS Biol 2006, 4:e291.

37. Lapraz F, Besnardeau L, Lepage T: Patterning of the dorsal-ventral axis in echinoderms: insights into the evolution of the BMP-chordin signaling network. PLoS Biol 2009, 7:e1000248.

38. Matus DQ, Thomsen GH, Martindale MQ: Dorso/ventral genes are asymmetrically expressed and involved in germ-layer demarcation during cnidarian gastrulation. Curr Biol 2006, 16:499–505.

39. Adamska M, Degnan SM, Green KM, Adamski M, Craigie A, Larroux C, Degnan BM: Wnt and TGF-beta expression in the sponge Amphimedon queenslandica and the origin of metazoan embryonic patterning. PLoS One 2007, 2:e1031.

40. Pang K, Ryan JF, Baxevanis AD, Martindale MQ: Evolution of the TGF-β signaling pathway and its potential role in the ctenophore, Mnemiopsis leidyi. PLoS One 2011, 6:e24152.

41. Watanabe H, Schmidt H, Kuhn A, Hoger S, Kocagoz Y, Laumann-lipp N, Suat O, Holstein TW: Nodal signalling determines biradial asymmetry. Nature 2014, 515:112–115.

42. Genikhovich G, Technau U: On the evolution of bilaterality. Development 2017, 144:3392–3404.

43. Garcia Abreu J, Coffinier C, Larraín J, Oelgeschläger M, De Robertis EM: Chordin-like CR domains and the regulation of evolutionarily conserved extracellular signaling systems. Gene 2002, 287:39–47.

44. Matsui M, Mizuseki K, Nakatani J, Nakanishi S, Sasai Y: Xenopus kielin: A dorsalizing factor containing multiple chordin-type repeats secreted from the embryonic midline. Proc Natl Acad Sci U S A 2000, 97:5291–6.

45. Srivastava M, Simakov O, Chapman J, Fahey B, Gauthier MEA, Mitros T, Richards GS, Conaco C, Dacre M, Hellsten U, Larroux C, Putnam NH, Stanke M, Adamska M, Darling A, Degnan SM, Oakley TH, Plachetzki DC, Zhai Y, Adamski M, Calcino A, Cummins SF, Goodstein DM, Harris C, Jackson DJ, Leys SP, Shu S, Woodcroft BJ, Vervoort M, Kosik KS, et al.: The Amphimedon queenslandica genome and the evolution of animal complexity. Nature 2010, 466:720–6.

46. Rentzsch F, Guder C, Vocke D, Hobmayer B, Holstein TW: An ancient chordin-like gene in organizer formation of Hydra. Proc Natl Acad Sci U S A 2007, 104:3249–54.

47. Putnam NH, Srivastava M, Hellsten U, Dirks B, Chapman J, Salamov A, Terry A, Shapiro H, Lindquist E, Kapitonov V V, Jurka J, Genikhovich G, Grigoriev I V, Lucas SM, Steele RE, Finnerty JR, Technau U, Martindale MQ, Rokhsar DS: Sea anemone genome reveals ancestral eumetazoan gene repertoire and genomic organization. Science 2007, 317:86–94.

48. Hui JHL, Holland PWH, Ferrier DEK: Do cnidarians have a ParaHox cluster? Analysis of synteny around a Nematostella homeobox gene cluster. Evol Dev 2008, 10:725–730.

49. DuBuc TQ, Ryan JF, Shinzato C, Satoh N, Martindale MQ: Coral comparative genomics reveal expanded Hox cluster in the cnidarian-bilaterian ancestor. Integr Comp Biol 2012, 52:835–41.

50. Simakov O, Marletaz F, Cho S-J, Edsinger-Gonzales E, Havlak P, Hellsten U, Kuo D-H, Larsson T, Lv J, Arendt D, Savage R, Osoegawa K, de Jong P, Grimwood J, Chapman JA, Shapiro H, Aerts A, Otillar RP, Terry AY, Boore JL, Grigoriev I V, Lindberg DR, Seaver EC, Weisblat DA, Putnam NH, Rokhsar DS: Insights into bilaterian evolution from three spiralian genomes. Nature 2013, 493:526–31.

51. Dattoli AA, Hink MA, DuBuc TQ, Teunisse BJ, Goedhart J, Röttinger E, Postma M: Domain analysis of the Nematostella vectensis SNAIL ortholog reveals unique nucleolar localization that depends on the zinc-finger domains. Sci Rep 2015, 5:12147.

52. Essex LJ, Mayor R, Sargent MG: Expression of Xenopus snail in mesoderm and prospective neural fold ectoderm. Dev Dyn 1993, 198:108–22.

53. Carver E a, Jiang R, Lan Y, Oram KF, Gridley T, Lan YU: The Mouse Snail Gene Encodes a Key Regulator of the Epithelial-Mesenchymal Transition The Mouse Snail Gene Encodes a Key Regulator of the Epithelial-Mesenchymal Transition. Mol Cell Biol 2001, 21:8184–8188.

54. Martindale MQ, Pang K, Finnerty JR: Investigating the origins of triploblasty: “mesodermal” gene expression in a diploblastic animal, the sea anemone Nematostella vectensis (phylum, Cnidaria; class, Anthozoa). Development 2004, 131:2463–2474.

55. Fritzenwanker JH, Saina M, Technau U: Analysis of forkhead and snail expression reveals epithelial-mesenchymal transitions during embryonic and larval development of Nematostella vectensis. Dev Biol 2004, 275:389–402.

56. Magie CR, Daly M, Martindale MQ: Gastrulation in the cnidarian Nematostella vectensis occurs via invagination not ingression. Dev Biol 2007, 305:483–497.

57. Marlow HQ, Srivastava M, Matus DQ, Rokhsar D, Martindale MQ: Anatomy and development of the nervous system of Nematostella vectensis, an anthozoan cnidarian. Dev Neurobiol 2009, 69:235–54.

58. Kelava I, Rentzsch F, Technau U: Evolution of eumetazoan nervous systems: insights from cnidarians. 2015(ii):1–24.

59. Rassat JJJ, Ruthmann A: Trichoplax adhaerens F.E. Schulze (placozoa) in the scanning electron microscope. Zoomorphologie 1979, 93:59–72.

60. Grell KG, Benwitz G: Additional investigations on the ultrastructure of Trichoplax adhaerens F.E. Schulze (Placozoa). Zoomorphology 1981, 98:47–67.

61. Ruthmann A, Behrendt GWR: The ventral epithelium of Trichoplax adhaerens (Placozoa): Cytoskeletal structures, teil contacts and endocytosis. Zoomorphology 1986, 106:115–122.

62. Behrendt G, Ruthmann A: The cytoskeleton of the fiber cells of Trichoplax adhaerens (Placozoa). Zoomorphology 1986:123–130.

63. Thiemann M, Ruthmann A: Zoomorphology Spherical forms of Trichoplax adhaerens (Placozoa). Zoomorphology 1990, 110:37–45.

64. Buchholz K, Ruthmann A: The Mesenchyme-Like Layer of the Fiber Cells of Trichoplax adhaerens (Placozoa), a Syncytium. Z Naturforsch 1995, 50c:282–285.

65. Spring J, Yanze N, Jösch C, Middel AM, Winninger B, Schmid V: Conservation of Brachyury, Mef2, and Snail in the Myogenic Lineage of Jellyfish: A Connection to the Mesoderm of Bilateria. Dev Biol 2002, 244:372–384.

66. Matus DQ, Pang K, Marlow H, Dunn CW, Thomsen GH, Martindale MQ: Molecular evidence for deep evolutionary roots of bilaterality in animal development. Proc Natl Acad Sci U S A 2006, 103:11195–11200.

67. Saina M, Genikhovich G, Renfer E, Technau U: BMPs and chordin regulate patterning of the directive axis in a sea anemone. Proc Natl Acad Sci U S A 2009, 106:18592–18597.

68. Genikhovich G, Fried P, Prünster MM, Schinko JB, Gilles AF, Fredman D, Meier K, Iber D, Technau U: Axis Patterning by BMPs: Cnidarian Network Reveals Evolutionary Constraints. Cell Rep 2015, 10:1646–1654.

69. Wijesena N, Simmons DK, Martindale MQ: Antagonistic BMP-cWNT signaling in the cnidarian Nematostella vectensis reveals insight into the evolution of mesoderm. PNAS 2017:E5608–E5615.

70. Finnerty JR, Pang K, Burton P, Paulson D, Martindale MQ: Origins of bilateral symmetry: Hox and dpp expression in a sea anemone. Science 2004, 304:1335–7.

71. Rentzsch F, Anton R, Saina M, Hammerschmidt M, Holstein TW, Technau U: Asymmetric expression of the BMP antagonists chordin and gremlin in the sea anemone Nematostella vectensis: implications for the evolution of axial patterning. Dev Biol 2006, 296:375–87.

72. Röttinger E, Dahlin P, Martindale MQ: A framework for the establishment of a cnidarian gene regulatory network for “endomesoderm” specification: the inputs of β-catenin/TCF signaling. PLoS Genet 2012, 8:e1003164.

73. Leclere L, Rentzsch F: RGM Regulates BMP-Mediated Secondary Axis Formation in the Sea Anemone Nematostella vectensis. Cell Rep 2014, 9:1921–1930.

74. Kraus Y, Aman A, Technau U, Genikhovich G: Pre-bilaterian origin of the blastoporal axial organizer. Nat Commun 2016, 7(May):11694.

75. Hayward DC, Samuel G, Pontynen PC, Catmull J, Saint R, Miller DJ, Ball EE: Localized expression of a dpp/BMP2/4 ortholog in a coral embryo. Proc Natl Acad Sci U S A 2002, 99:8106–8111.

76. Reber-Müller S, Streitwolf-Engel R, Yanze N, Schmid V, Stierwald M, Erb M, Seipel K: BMP2/4 and BMP5-8 in jellyfish development and transdifferentiation. Int J Dev Biol 2006, 50:377–84.

77. Yamada A, Pang K, Martindale MQ, Tochinai S: Surprisingly complex T-box gene complement in diploblastic metazoans. Evol Dev 2007, 9:220–30.

78. Ryan JF, Pang K, Mullikin JC, Martindale MQ, Baxevanis AD: The homeodomain complement of the ctenophore Mnemiopsis leidyi suggests that Ctenophora and Porifera diverged prior to the ParaHoxozoa. Evodevo 2010, 1:9.

79. Hayward DC, Catmull J, Reece-Hoyes JS, Berghammer H, Dodd H, Hann SJ, Miller DJ, Ball EE: Gene structure and larval expression of cnox-2Am from the coral Acropora millepora. Dev Genes Evol 2001, 211:10–19.

80. Ryan JF, Mazza ME, Pang K, Matus DQ, Baxevanis AD, Martindale MQ, Finnerty JR: Pre-bilaterian origins of the hox cluster and the hox code: Evidence from the sea anemone, Nematostella vectensis. PLoS One 2007, 2:e153.

81. Miljkovic-Licina M, Chera S, Ghila L, Galliot B: Head regeneration in wild-type hydra requires de novo neurogenesis. Development 2007, 134:1191–1201.

82. Arnone MI, Rizzo F, Annunciata R, Cameron RA, Peterson KJ, Martínez P: Genetic organization and embryonic expression of the ParaHox genes in the sea urchin S. purpuratus: insights into the relationship between clustering and colinearity. Dev Biol 2006, 300:63–73.

83. Wollesen T, Rodríguez Monje SV, McDougall C, Degnan BM, Wanninger A: The ParaHox gene Gsx patterns the apical organ and central nervous system but not the foregut in scaphopod and cephalopod mollusks. Evodevo 2015, 6:41.

84. Illes JC, Winterbottom E, Isaacs H V: Cloning and expression analysis of the anterior parahox genes, Gsh1 and Gsh2 from Xenopus tropicalis. Dev Dyn 2009, 238:194–203.

85. Pei Z, Wang B, Chen G, Nagao M, Nakafuku M, Campbell K: Homeobox genes Gsx1 and Gsx2 differentially regulate telencephalic progenitor maturation. Proc Natl Acad Sci U S A 2011, 108:1675–80.

86. Ikuta T, Chen Y-C, Annunziata R, Ting H-C, Tung C, Koyanagi R, Tagawa K, Humphreys T, Fujiyama A, Saiga H, Satoh N, Yu J-K, Arnone MI, Su Y-H: Identification of an intact ParaHox cluster with temporal colinearity but altered spatial colinearity in the hemichordate Ptychodera flava. BMC Evol Biol 2013, 13:129.

87. Garstang MG, Osborne PW, Ferrier DEK: TCF/Lef regulates the Gsx ParaHox gene in central nervous system development in chordates. BMC Evol Biol 2016, 16:57.

88. Nakanishi N, Renfer E, Technau U, Rentzsch F: Nervous systems of the sea anemone Nematostella vectensis are generated by ectoderm and endoderm and shaped by distinct mechanisms. Development 2012, 139:347–57.

89. Cambell R: Elimination of Hydra interstitial and nerve cells by means of colchicine. J Cell Sci 1976, 21:1–13.

90. Fujisawa T: Role of interstitial cell migration in generating position-dependent patterns of nerve cell differentiation in Hydra. Dev. Dev Biol 1989, 133:77–82.

91. Künzel T, Heiermann R, Frank U, Müller W, Tilmann W, Bause M, Nonn A, Helling M, Schwarz RS, Plickert G: Migration and differentiation potential of stem cells in the cnidarian Hydractinia analysed in eGFP-transgenic animals and chimeras. Dev Biol 2010, 348:120–129.

92. Martin V, Thomas M: The origin of the nervous system in Pennaria tiarella, as revealed by treatment with colchicine. Biol Bull 1981, 160:303–310.

93. Thomas M, Freeman G, Martin V: The embryonic origin of neurosensory cells and the role of nerve cells in metamorphosis in Phiolidium gregarium (Cnidaria, Hydrozoa). Int J Invert Reprod Dev 1987, 11:265–285.

94. Senatore A, Reese TS, Smith CL: Neuropeptidergic integration of behavior in Trichoplax adhaerens, an animal without synapses. J Exp Biol 2017, 220:3381–3390.

95. Smith CL, Abdallah S, Wong YY, Le P, Harracksingh AN, Artinian L, Tamvacakis AN, Rehder V, Reese TS, Senatore A: Evolutionary insights into T-type Ca 2 + channel structure, function, and ion selectivity from the Trichoplax adhaerens homologue. J Gen Physiol 2017:1–28.

96. Furuya H, Tsuneki K: Biology of dicyemid mesozoans. Zoolog Sci 2003, 20:519–532.

97. Suzuki TG, Ogino K, Tsuneki K, Furuya H: Phylogenetic analysis of dicyemid mesozoans (phylum Dicyemida) from innexin amino acid sequences: dicyemids are not related to Platyhelminthes. J Parasitol 2010, 96:614–625.

98. Ryan JF, Pang K, Schnitzler CE, Nguyen A-D, Moreland RT, Simmons DK, Koch BJ, Francis WR, Havlak P, Smith S a, Putnam NH, Haddock SHD, Dunn CW, Wolfsberg TG, Mullikin JC, Martindale MQ, Baxevanis AD: The genome of the ctenophore Mnemiopsis leidyi and its implications for cell type evolution. Science 2013, 342:1242592.

99. Shinzato C, Shoguchi E, Kawashima T, Hamada M, Hisata K, Tanaka M, Fujie M, Fujiwara M, Koyanagi R, Ikuta T, Fujiyama A, Miller DJ, Satoh N: Using the Acropora digitifera genome to understand coral responses to environmental change. Nature 2011, 476:320–323.

100. Colbourne JK, Pfrender ME, Gilbert D, Thomas WK, Tucker A, Oakley TH, Tokishita S, Aerts A, Arnold GJ, Basu MK, Bauer DJ, Cáceres CE, Carmel L, Casola C, Choi J-H, Detter JC, Dong Q, Dusheyko S, Eads BD, Fröhlich T, Geiler-Samerotte KA, Gerlach D, Hatcher P, Jogdeo S, Krijgsveld J, Kriventseva E V, Kültz D, Laforsch C, Lindquist E, Lopez J, et al.: The ecoresponsive genome of Daphnia pulex. Science 2011, 331:555–561.

101. Putnam NH, Butts T, Ferrier DEK, Furlong RF, Hellsten U, Kawashima T, Robinson-Rechavi M, Shoguchi E, Terry A, Yu J-K, Benito-Gutiérrez EL, Dubchak I, Garcia-Fernàndez J, Gibson-Brown JJ, Grigoriev I V, Horton AC, de Jong PJ, Jurka J, Kapitonov V V, Kohara Y, Kuroki Y, Lindquist E, Lucas S, Osoegawa K, Pennacchio LA, Salamov AA, Satou Y, Sauka-Spengler T, Schmutz J, Shin-I T, et al.: The amphioxus genome and the evolution of the chordate karyotype. Nature 2008, 453:1064–1071.

102. Schultz J, Milpetz F: SMART, a simple modular architecture research tool: identification of signaling domains. Proc … 1998, 95(May):5857–5864.

103. Edgar RC: MUSCLE: multiple sequence alignment with high accuracy and high throughput. Nucleic Acids Res 2004, 32:1792–7.

104. Huelsenbeck JP, Ronquist F: MRBAYES: Bayesian inference of phylogeny. Bioinformatics 2001, 17:754–755.

105. Ronquist F, Huelsenbeck JP: MrBayes 3: Bayesian phylogenetic inference under mixed models. Bioinformatics 2003, 19:1572–1574.

106. FigTree [http://tree.bio.ed.ac.uk/software/figtree]

